# Nucleotide sequence and DNaseI sensitivity are predictive of 3D chromatin architecture

**DOI:** 10.1101/103614

**Authors:** Jacob Schreiber, Maxwell Libbrecht, Jeffrey Bilmes, William Stafford Noble

**Affiliations:** Paul G. Allen School of Computer Science, University of Washington, Seattle, USA; Department of Genome Sciences, University of Washington, Seattle, USA; Department of Electrical Engineering, University of Washington, Seattle, USA

## Abstract

Recently, Hi-C has been used to probe the 3D chromatin architecture of multiple organisms and cell types. The resulting collections of pairwise contacts across the genome have connected chromatin architecture to many cellular phenomena, including replication timing and gene regulation. However, high resolution (10 kb or finer) contact maps remain scarce due to the expense and time required for collection. A computational method for predicting pairwise contacts without the need to run a Hi-C experiment would be invaluable in understanding the role that 3D chromatin architecture plays in genome biology. We describe Rambutan, a deep convolutional neural network that predicts Hi-C contacts at 1 kb resolution using nucleotide sequence and DNaseI assay signal as inputs. Specifically, Rambutan identifies locus pairs that engage in high confidence contacts according to Fit-Hi-C, a previously described method for assigning statistical confidence estimates to Hi-C contacts. We first demonstrate Rambutan’s performance across chromosomes at 1 kb resolution in the GM12878 cell line. Subsequently, we measure Rambutan’s performance across six cell types. In this setting, the model achieves an area under the receiver operating characteristic curve between 0.7662 and 0.8246 and an area under the precision-recall curve between 0.3737 and 0.9008. We further demonstrate that the predicted contacts exhibit expected trends relative to histone modification ChlP-seq data, replication timing measurements, and annotations of functional elements such as promoters and enhancers. Finally, we predict Hi-C contacts for 53 human cell types and show that the predictions cluster by cellular function. [NOTE: After our original submission we discovered an error in our calling of statistically significant contacts. Briefly, when calculating the prior probability of a contact, we used the number of contacts at a certain genomic distance in a chromosome but divided by the total number of bins in the full genome. When we corrected this mistake we noticed that the Rambutan model, as it curently stands, did not outperform simply using the GM12878 contact map that Rambutan was trained on as the predictor in other cell types. While we investigate these new results, we ask that readers treat this manuscript skeptically.]

## 1 Introduction

3D chromatin architecture is known to be involved in many cellular phenomena including, but not limited to, gene regulation, replication timing, and disease. Currently, large-scale chromatin architecture data is collected through Hi-C (Lieberman-Aiden *et al*., 2009), ChIA-PET (Fullwood *et al*., 2009), or Hi-ChIP (Mumbach *et al*., 2016) experiments that identify, on a genome-wide basis, which fragments of the human genome are in contact. These procedures, while powerful, are both time intensive and cost prohibitive to perform at high resolution (10 kb or finer). The highest resolution contact map produced thus far has been a 1 kb resolution contact map of GM12878 which required 4.9 billion paired-end reads to generate (Rao *et al*., 2014).

This scarcity of high resolution experimental data is problematic because differences in chromatin architecture across cell types from the same organism may have significant impact on processes such as gene regulation and DNA replication (Lieberman-Aiden *et al*., 2009; Rao *et al*., 2014; Ma *et al*., 2015; Heidari *et al*., 2014). This variation implies that nucleotide sequence information alone cannot serve as the sole input to a predictive model of chromatin architecture. Such a model will require data from a cell type specific assay, preferably one that is inexpensive and has already been applied to a wide variety of organisms and cell types. A good candidate assay is DNaseI sensitivity (John *et al*., 2013), because this data is cheap to collect and has already been obtained for hundreds of cell types, including 443 human cell types in the ENCODE project as of December 2016. Hi-C is more expensive than other genomics assays such as DNaseI because, to achieve a given level of precision, it requires a number of sequencing reads that grows quadratically with the length of the genome. We propose a model that predicts Hi-C contact maps using as input only nucleotide sequence information and DNaseI sensitivity.

Although the primary output of a Hi-C experiment consists of counts of contacts between pairs of loci, we train our model to predict statistical confidence estimates associated with those contacts, rather than the raw counts. We chose this target for our modeling effort because the raw contact counts include biases that reflect physical and experiment artifacts rather than interesting biology. Most notably, the counts are dominated by the genomic distance effect, i.e., by contacts among pairs of proximal loci that come into contact due to random self-looping of the DNA polymer. In addition, Hi-C data is known to include biases reflective of local GC content and mappability (Imakaev *et al*., 2012). Methods such as Fit-Hi-C (Ay *et al*., 2014) assign statistical confidence estimates to contacts while accounting for these types of biases. Thus, we pose the prediction problem as a classification task which predicts whether a given pair of loci are in contact with high confidence, as determined by Fit-Hi-C. We also chose to predict fine-scale contacts rather than large-scale features of the structure of the genome, such as topologically associated domains. Nonetheless, as we demonstrate below, the predictions made by our model successfully recapitulate many large-scale features of Hi-C data.

In addition to choosing inputs and outputs, we also had to decide at what resolution we should attack this prediction problem. In general, the appropriate resolution for analyzing a Hi-C contact map depends on the sequencing depth. In practice, publicly available Hi-C data sets vary widely in their effective resolution. The lymphoblastoid cell line GM12878 is the only cell type that has been sequenced deeply enough to produce a 1 kb map, though K562, IMR90, NHEK, HMEC, and HUVEC have all been sequenced deeply enough to produce 5 kb maps (Rao *et al*., 2014). We decided to train and test the model in two settings. First, we train a 1 kb model on some GM12878 chromosomes and validate on the held-out chromosomes. Second, we train on all GM12878 chromosomes and validate on Hi-C data from other cell types. In the latter case, we convert our 1 kb resolution predictions to 5 kb resolution in order to compare to 5 kb resolution data in other cell types (Section 3).

For the predictive model, we employ a deep multimodal convolutional neural network. Deep learning has recently been applied successfully to several important biological problems (Kelley *et al*., 2016; Zhou and Troyanskaya, 2015; Wang *et al*., 2017; Leung *et al*., 2014). Such methods are attractive due to their ability to extract complex yet meaningful feature representations from massive data sets without requiring extensive feature engineering. To our knowledge, machine learning methods have not previously been used to predict Hi-C contacts in a systematic fashion, though Fortin and Hansen (2015) have shown how to predict large-scale Hi-C structure from histone modification ChIP-seq data and Whalen *et al*. (2016) predict specific enhancer-promoter interactions. In the cross-chromosome setting in GM12878, our model, called Rambutan, achieves an area under the receiver operating characteristic (AUROC) curve of 0.846, and a corresponding area under the precision-recall (AUPR) curve of 0.373. In the cross-cell type setting, the model achieves between a 0.766 and 0.825 AUROC and between a 0.374 and 0.901 AUPR, outperforming both a ranking based on genomic distance and using the contact map for GM12878 as a predictor for all cell types.

To further validate Rambutan’s performance, we investigated a variety of other expected properties of Rambutan’s predictions. The insulation score quantifies how patterns of local contacts resemble the pattern expected near a domain boundary. This score has been used in calling topologically associating domains (TADs) (Crane *et al*., 2015). We show that insulation scores calculated based on Rambutan’s predictions correlate strongly with insulation scores calculated from real Hi-C contact maps. Furthermore, both types of insulation scores correlate strongly with several histone modifications and with replication timing. In addition, Rambutan’s predictions confirm prior work showing that genomic loci linked by high confidence contacts tend to exhibit similar replication timing. Rambutan’s predictions show an expected enrichment in promoter and enhancer elements across multiple cell types, and in general Rambutan’s enrichment pattern across functional elements agrees with that of contacts called by Fit-Hi-C. Lastly, we use Rambutan to predict contact maps for chromosome 21 in 53 cell types, using data from the Roadmap Epigenomics project. These results show that cell types of similar function share similar structures, whereas cancer cell types are very different from each other.

## 2 Approach

In general, neural networks are extremely modular in their choice of layer composition, loss function, optimizer, regularization, and hyperparameters for the above. We describe each of these components briefly and suggest Schmidhuber (2015) as a comprehensive review.

**Model input and output** Rambutan takes nucleotide sequence and DNase sensitivity for two 1 kb loci as inputs and returns the probability that the pair engages in a high confidence contact (Fig. 1a). The labels used for training are binary where the positive label indicates the locus pair has a q-value ≤ 1e-6 and the negative label corresponds to a q-value above that threshold. Nucleotide sequence is one-hot encoded, providing 4 bits per position but activating only the one corresponding to the nucleotide present. DNase sensitivity is represented as log fold change over a control. For input into the model, this value is rounded to the nearest integer and used to activate the bits “up or down” to that number ranging from −2 to 5. For example, a log fold change value of 2.3 would be rounded to 2, activating bits 0 through 2, and a log fold change of −1.3 would be rounded to −1 and activate bits 0 and −1. This strategy allows the network to learn the marginal effect that differing values of DNase signal have. Lastly, genomic distance is encoded as 95 bits, each representing a different distance threshold, with all bits activated up until the threshold is reached. The first 50 bits represent marginal increases of 1 kb ranging from 50 kb to 100 kb, the next 40 bits represent marginal increases of 10 kb ranging from 100 kb to 500 kb, and the last five bits represent marginal increases of 100 kb ranging from 500 kb to 1 Mb. Thus, the network receives as input a total of 24,095 bits for each pair of loci. Empirically, using entirely binary inputs improved performance compared to a network in which some inputs were binary and others were continuous valued.

**Figure 1:**
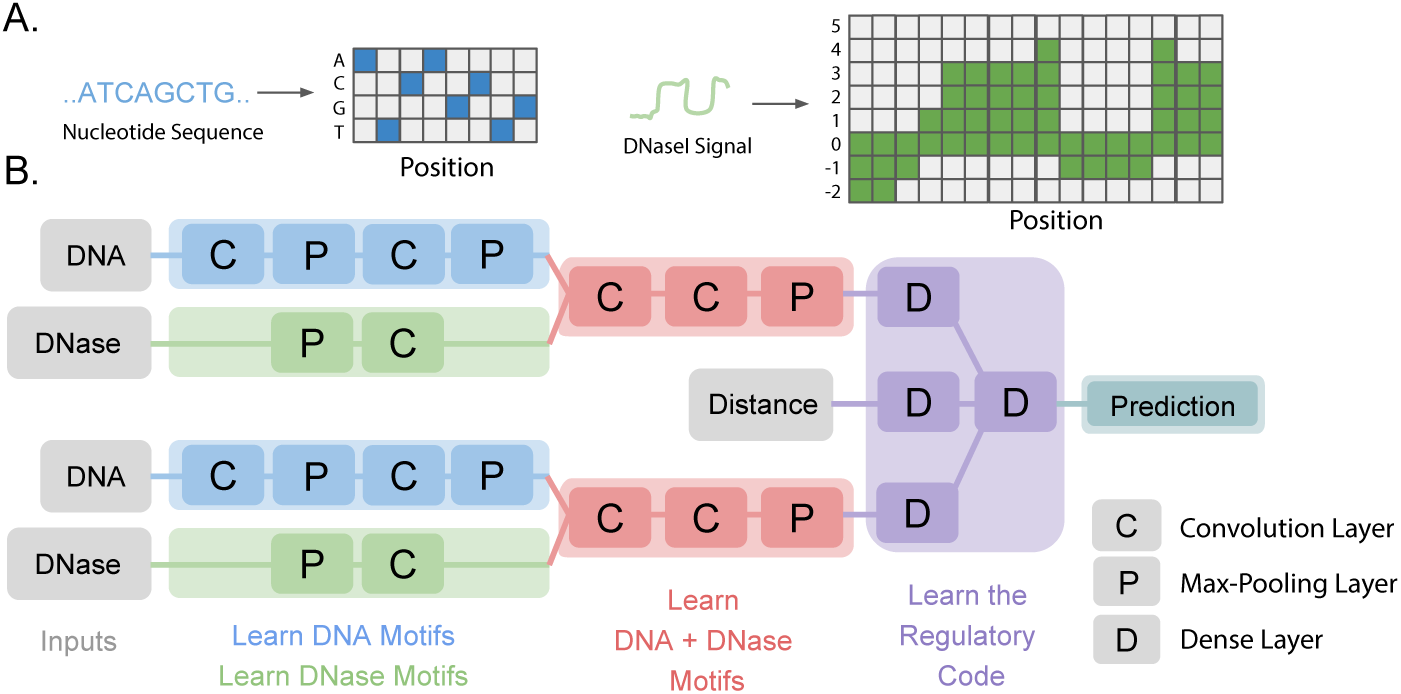
The inputs to and architecture of the Rambutan network. (a) Schematic showing how nucleotide sequence and DNase fold-change are encoded for input into the network. (b) In Rambutan, inputs flow from left to right, starting with four sources of input data and ending with a final prediction. Each symbol represents a layer in the network, colored by type: a “C” represents a convolutional layer, a “P” represents a max pooling layer, and a “D” represents a dense inner product layer. All layers are followed by a batch normalization layer then a ReLU activation, except for the final prediction layer which is just a two-node dense layer with a softmax activation for binary prediction. The blue symbols indicate layers that learn DNA-specific patterns, the green symbols indicate layers that learn DNase-specific patterns, red symbols represent layers that learn patterns comprised of both DNA and a DNase components, and purple layers learn a form of regulatory code that interprets how these patterns are predictive of a contact. Feature maps learned from the blue and green layers are concatenated before being fed into the red layers.

**Network topology** In the Rambutan model, convolution and max-pooling layers are first used to extract meaningful features from the inputs and then dense inner-product layers are used to learn a regulatory code relating these features to high confidence contacts (Fig. 1b). In the context of sequence analysis, convolutions can be thought of as position weight matrices which are scanned along input sequences and output the affinity for that motif at that position. Each convolution and dense inner-product layer in the network is followed by batch normalization (Ioffe and Szegedy, 2015) and uses a rectified linear (ReLU) non-linearity. Rambutan has two arms that process the data from the two loci independently. Each of these locus-specific arms is comprised of two subarms that process, respectively, nucleotide sequence and DNase signal. These locus-specific arms then combine information from those signals to identify patterns corresponding to identify nucleotide and DNaseI patterns. More details on the topology is available in Suppl. Note 1.

**Training** Rambutan was trained using standard techniques. We chose to use the ADAM optimizer, given its recent success in the field of computer vision (Kingma and Ba, 2015), and the logistic loss function (Buja *et al*., 2005). All hyperparameters for both the ADAM optimizer and batch normalization layers were set to the defaults suggested by their respective papers and are specified in Suppl. Note 2. No weight decay or other form regularization, such as dropout, was used because empirically these regularizers did not affect validation performance. Training was performed on balanced batches of data until convergence was observed. Details can be found in Suppl. Note 2.

## 3 Methods

**Datasets** The datasets used in the experiment are fully described in Suppl. Note 3. Briefly, nucleotide sequence is from the hg19 reference genome, epigenetic data is from the Roadmap Epigenomics Consortium, replication timing data is from www.replicationdomain.org, and the Hi-C data is from Rao *et al*. (2014).

**Changing resolution** Rambutan makes predictions at 1 kb resolution that can be processed to obtain 5 kb resolution contact maps. Each pair of 5 kb loci corresponds to 25 pairs of 1 kb loci. The procedure involves using either the maximum Rambutan prediction or the minimum Fit-Hi-C p-value among those 25, depending on which type of data is being analyzed. The max function treats the values within that square as not being independent from each other, and was found empirically to perform the best among several other methods.

**Insulation Score** The insulation score measures the extent to which a given genomic locus exhibits a pattern of local contacts indicative of the boundary of a TAD (Crane *et al*., 2015). Calculating the score involves running a square down the diagonal of the contact map and summing the number of contacts within that square for each position in the chromosome. This score is then converted into a log fold enrichment over the average number of contacts:

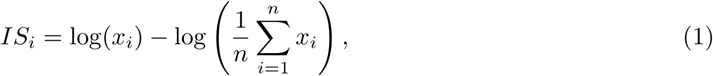

where *x_i_* is the sum of contacts within the square surrounding locus *i*, and *IS_i_* is the insulation score of locus *i*. We calculate the insulation score using a 1 Mb square for both the original Hi-C maps and for the Rambutan predictions. The latter calculation sums the probability of each pair of loci being in contact in lieu of the number of contacts.

## 4 Results

### 4.1 Rambutan makes accurate 1 kb resolution predictions

We first tested how well Rambutan can make 1 kb resolution predictions. Because GM12878 is the only cell type for which a 1 kb resolution resolution contact map is available, we performed a cross-chromosome validation: we trained Rambutan on chromosomes 1-20, validated the model on chromosome 21, and tested its final performance on chromosome 22. In this experiment, Rambutan achieved a test set AUROC of 0.846 and an AUPR of 0.373 (Fig. 2). For comparison, simpler models such as logistic regression and a random forest classifier performed no better than using genomic distance alone.

**Figure 2:**
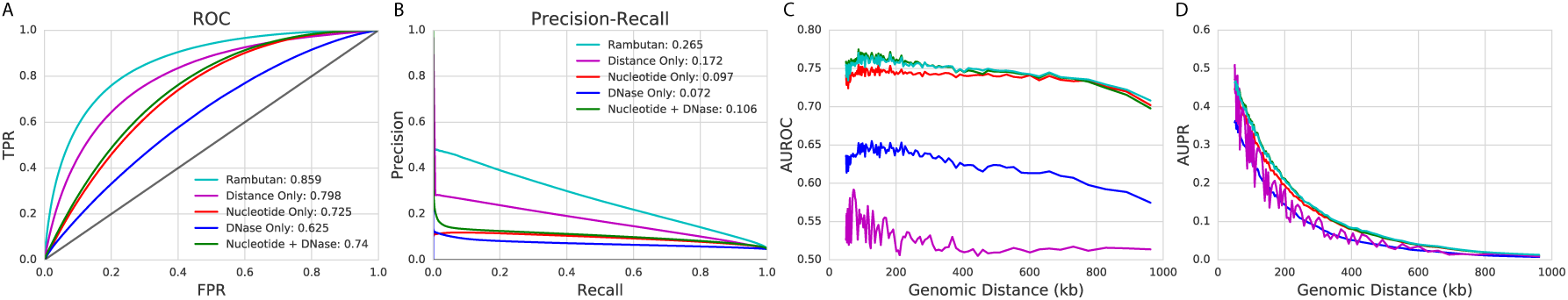
Performance of the Rambutan model at 1 kb resolution. (a and b) ROC curves and PR curves for the full Rambutan model are shown and compared to those from other baselines. The area under each of these curves is shown in the legend. (c and d) The area under the ROC curve and area under the PR curve is shown as a function of genomic distance. Since the sparsity of contacts increases as a function of genomic distance the measurements at further distances would be very imprecise. We handle this by instead using percentiles such that each point contains 1% of all true contacts when ordered by genomic distance. All results are shown on the validation set.

Given that the model made accurate predictions, we then tested the effect that each input had on the performance of the model on the validation set to ensure that it was leveraging both nucleotide sequence and DNase signal. The full model was compared to a ranking based on genomic distance alone and to models using as inputs (1) only nucleotide sequence, (2) only DNase sensitivity, and (3) both nucleotide sequence and DNase sensitivity but not genomic distance (Fig. 2). The results confirm that nucleotide sequence and DNase can be leveraged together for better performance than using either input separately. Strikingly, despite the use of Fit-Hi-C to control for the genomic distance effect, simply ranking based on the distance between two loci yields very strong predictive performance. This is not unexpected, because the density of contacts near the diagonal of the matrix means that Fit-Hi-C has much better statistical power to detect contacts, even after controlling for random DNA looping. Of course, such a naive model would output fixed predictions at each genomic distance and be useless at making biologically relevant predictions. Overall, the full Rambutan model strongly outperforms each of the other methods that we investigated.

Because we saw a strong genomic distance effect, we wanted to confirm that Rambutan performed well at all genomic distances. We did this by evaluating model performance as a function of genomic distance (Fig. 2c-d). The results indicate that Rambutan performs well at all genomic distances, with AUROC not decreasing significantly as distance increases. We note that, in this setting, the full model performs similarly to using only nucleotide sequence and DNase signal (Fig. 2), which makes sense because we are factoring out distance as an important measurement.

### 4.2 Rambutan makes cell type specific predictions

Having established that Rambutan can make 1 kb resolution predictions accurately we turn to the more challenging, but more interesting, task of predicting contact maps for other cell types. Five other human cell types have been sequenced deeply enough to produce 5 kb resolution contact maps: K562, IMR90, NHEK, HMEC, and HUVEC. Using the model trained in GM12878, we predict 1 kb resolution contact maps for each of these cell types and then convert the signal to 5 kb resolution (Section 3). Because a large portion of 3D chromatin architecture is shared across cell types, we compare Rambutan to simply using the GM12878 contact map as a predictor for other cell types. The Fit-Hi-C p-values from the 1 kb resolution GM12878 contact map were converted to 5 kb resolution in the same manner as the Rambutan predictions to control for error introduced in this step. We also use genomic distance as a predictor, as before.

In cross-cell type predictions, Rambutan outperforms both baselines across almost all cell types, achieving AUROCs 0.7662-0.8246 and AUPRs of 0.3737-0.9008 (Figure 4). Results for all cell types are shown in Suppl. Note 4. One important observation is that using the GM12878 contact map as a predictor does not yield perfect performance in the GM12878 cell type because the process of changing the resolution introduces error in the Fit-Hi-C significance estimates. Likewise, the convex segments in the ROC curves can likely be explained both by the 1 kb-to-5 kb conversion and as an artifact introduced by aggregating Fit-Hi-C p-values across multiple distances.

An interesting observation is that, in the NHEK cell type, ranking pairs of loci by genomic distance (AUROC 0.8033) outperforms Rambutan predictions (AUROC 0.772) and GM12878 (AUROC 0.7161). NHEK shows an inconsistency across a variety of evaluation metrics (Fig. 3 and 5), suggesting either anomalous data or markedly different 3D architecture in that cell type. We note that across several evaluation metrics the Rambutan predictions remain consistant with known biology while the Hi-C derived insulation score is not consistent, suggesting technical error. In addition, among Hi-C derived insulation scores, NHEK is an outlier and does not correlate well with the others (Supp Fig. 5). Standard quality measures show no obvious problems with the Hi-C contact map from NHEK despite these inconsistencies.

**Figure 3:**
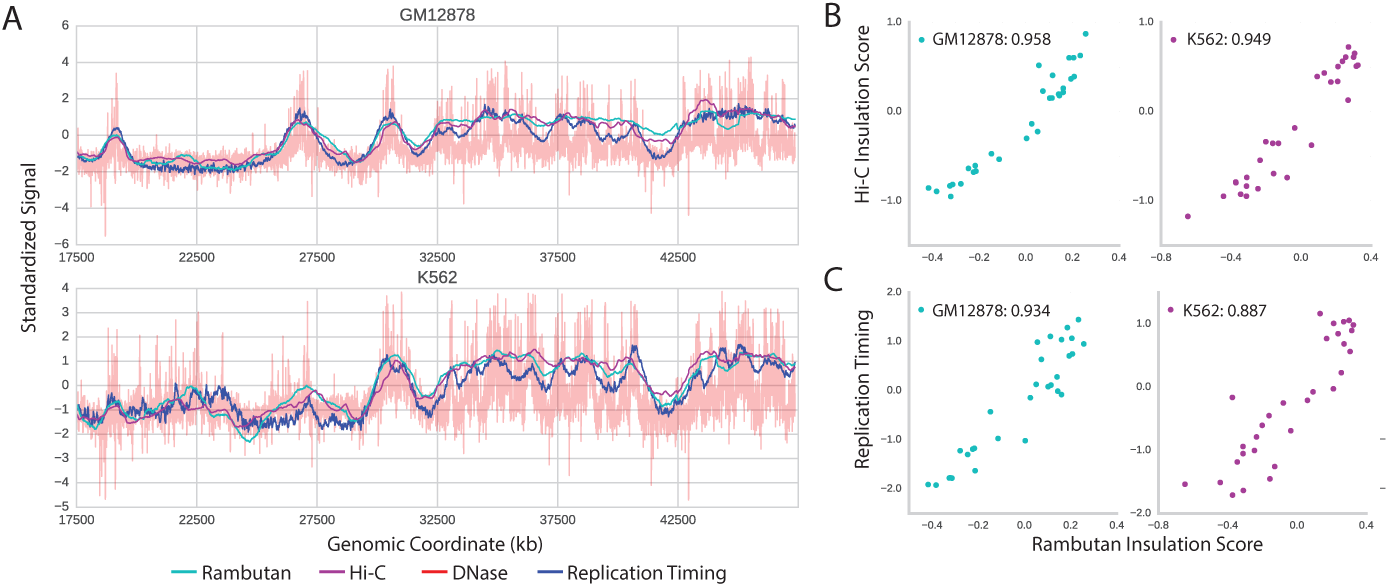
Comparison of insulation scores derived from real Hi-C data and Rambutan predictions. (a) The Rambutan derived insulation score (cyan) is compared to both the Hi-C derived insulation score (magenta) and DNase sensitivity (red). Two cell types are shown to highlight differences across all three measurements. The insulation scores are smooth because the score at each locus is highly correlated with the loci adjacent to it. All values have been standardized for display purposes. (b) The Rambutan derived insulation score is compared to the Hi-C derived insulation score at each independent measurement, and the Pearson correlation is calculated between the two. (c) The same insulation scores are shown compared to replication timing (blue). (d) The Ramutan derived insulation score is compared to replication timing at each independent measurement and the Pearson correlation is calculated between the two.

**Figure 4:**
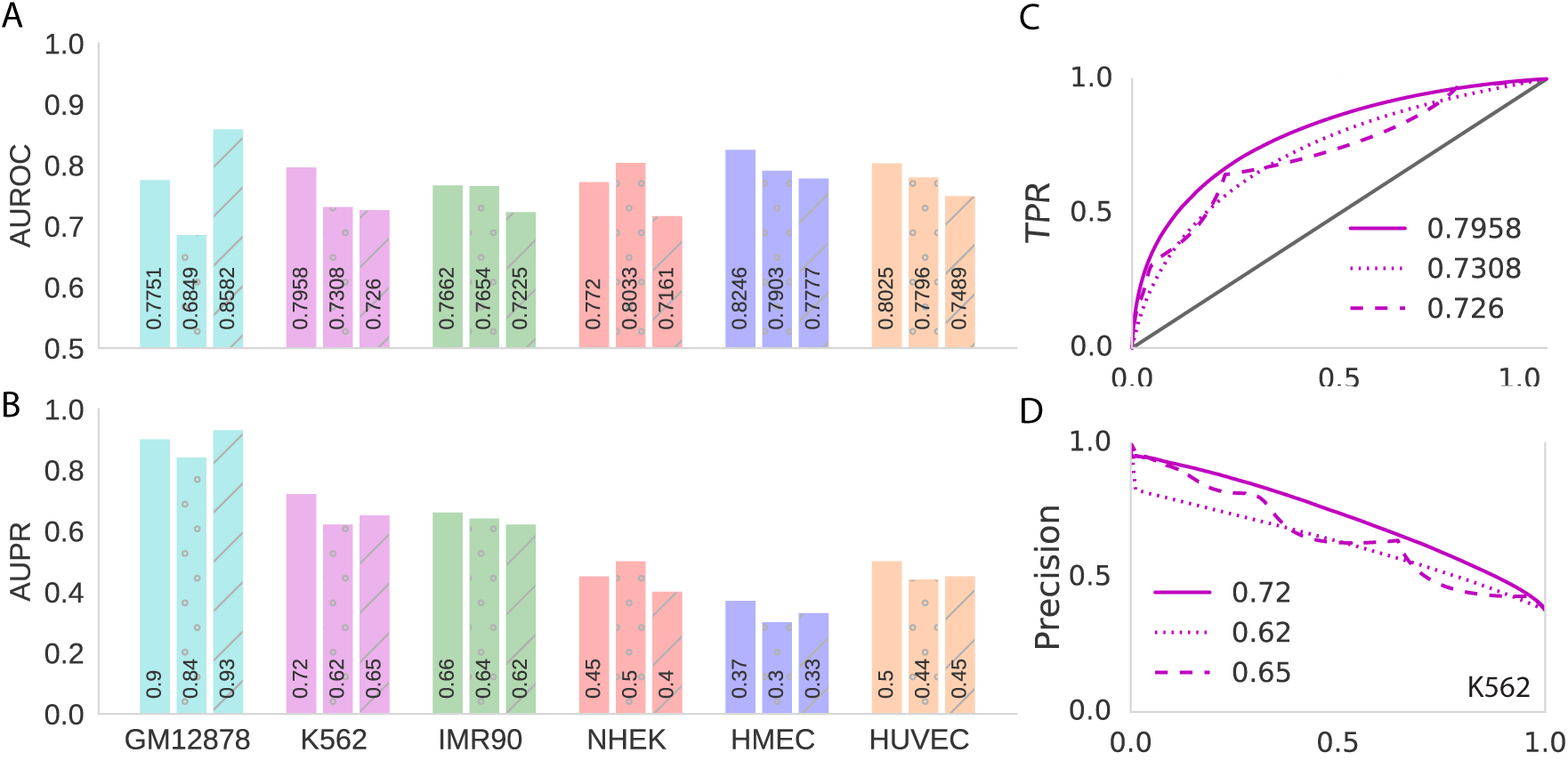
Performance of the Rambutan model across six cell types. (a and b) AUROC and AUPR measures when using Rambutan predictions (solid), genomic distance (dotted), or GM12878 (striped) as the predictor for all six cell types. Rambutan outperforms both baselines in all cell types except for genomic distance in NHEK. (c and d) Respectively, the ROC and PR curves for all three predictors in K562.

**Figure 5:**
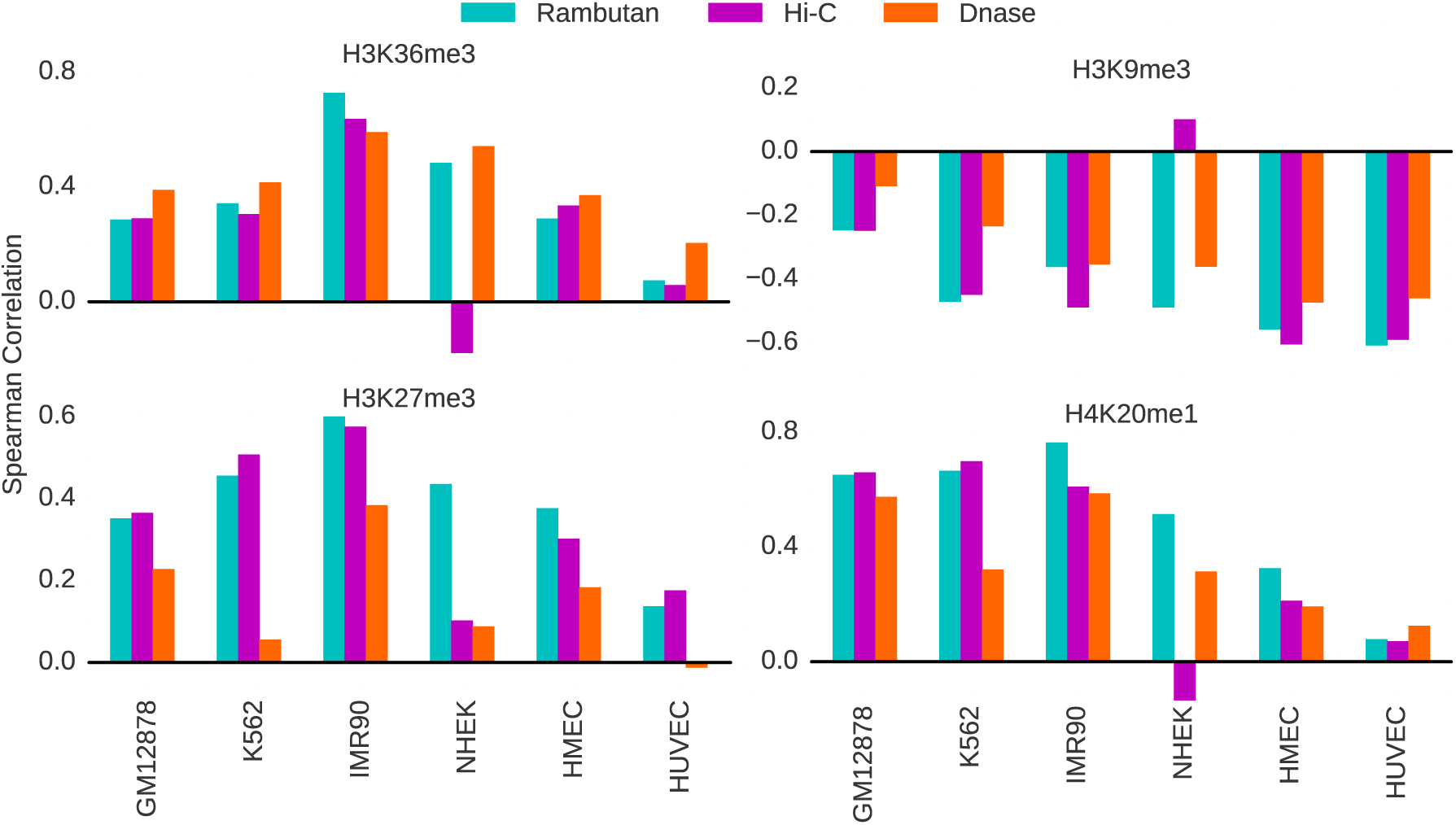
Correlation between insulation score and histone modifications. The Spearman correlation was calculated between the two insulation scores and four histone modification ChIP-seq values. The three histone marks that display peaks at regulatory elements—H3K36me3, H3K27me3, and H4K20me1—exhibit a corresponding positive correlation with insulation score, whereas the histone modification associated with constitutively silent domains, H3K9me3, exhibits a negative correlation with insulation score. These values are compared to the correlation between DNase signal and each histone modification, to control for the effect DNase may have as an input into the Rambutan model.

### 4.3 Rambutan predictions re-create the insulation score

Although performing well on metrics such as AUROC and AUPR are good indicators of success, we also wanted to be sure that the Rambutan predictions exhibit other properties associated with real Hi-C data. We therefore connected Rambutan’s predictions to the larger context of 3D chromatin architecture by showing that Rambutan predictions can identify domain-scale phenomena. The insulation score (Section 3) is designed to measure the extent to which Hi-C contacts near a given locus resemble the pattern expected near the boundary of two topologically associating domains (TADs). We chose to look at insulation scores rather than TAD calls because the insulation score is a continuous measurement, whereas TAD calls are discrete.

We calculated insulation scores from Hi-C contact maps and Rambutan predictions across all six cell types. Qualitatively, the shapes of the plots of real and predicted insulation scores at 1 Mb resolution match well, with coincident peaks and dips (Fig. 3a/b). The peaks in the insulation score exclusively appear in regions of DNase enrichment, as would be expected because both are measurements of chromatin accessibility. The Pearson correlations between the real and predicted insulation scores are high across all cell types, with most values approximately 0.95. A notable exception to this trend is the NHEK cell type whose experimental Hi-C data remains an outlier across all validation metrics. Similar overall trends hold when we re-calculate the insulation score at 100 kb or 40 kb rather than 1 Mb resolution (Supp Fig. 6 & 7), with correlations of 0.827-0.927 and 0.624-0.820, respectively, excluding NHEK.

Chromatin architecture has previously been connected to replication timing through eigenvector analysis of a processed Hi-C contact map (Dekker *et al*., 2013; Imakaev *et al*., 2012; Fortin and Hansen, 2015). Our next step was to connect Rambutan’s predictions to replication timing. Knowing that the eigenvector analysis will not work on a matrix of probabilities, we linked the predictions to replication timing through the insulation score. (Fig. 3c/d). Much like the comparison of two insulation scores, the Rambutan predicted insulation score matches the replication timing profile both in shape and with a strong Pearson correlation, between 0.723 (HUVEC) and 0.934 (GM12878).

Lastly, we wanted to connect Rambutan predicted insulation scores to histone modifications for four well-studied modifications: H3K36me3, H3K27me3, H4K20me1, and H3K9me3. Accordingly, we calculated the Spearman correlation between each insulation score and each histone modification ChIP-seq data set for each cell type (Fig. 5). DNase signal by itself is known to correlate with several histone modifications (Boyle *et al*., 2011), so we expected Rambutan’s predictions to share this correlation. However, the correlations derived from Rambutan’s predictions show far more similarity to those derived from the real insulation score than to those from DNase signal alone. Further supporting evidence of the relevance of this signal is that both sets of insulation scores positively correlate with histone marks that are associated with regulatory elements (H3K36me3, H3K27me3, and H4K20me1) but negatively correlate with a histone mark that occurs only at constitutively silent domains (H3K9me3). Prior work by Huang *et al*. (2015) supports this trend, showing that H3K9me3 is depleted near TAD boundaries, whereas the other histone marks are enriched near TAD boundaries. Again, NHEK is an exception, exhibiting the expected patterns of correlations with DNase and Rambutan predicted insulation scores, but opposite trends with real insulation scores.

### 4.4 Synchronously replicating regions are linked by high-confidence Rambutan calls

Several previous studies have established a connection between 3D chromatin architecture and replication timing. On a large scale, Ryba *et al*. (2010) showed a connection between the replication timing of two loci and their spatial proximity as measured by Hi-C, and Pope *et al*. (2014) demonstrated that boundaries in replication timing are indicative of boundaries between TADs. Ay *et al*. (2014) then built upon this work to show that loci linked by high-confidence Hi-C contacts share similar replication timing. Furthermore, Ay *et al*. (2014) suggest that this effect is most noticeable when comparing loci greater than ~750 kb apart in humans cell types.

We carried out a similar analysis with Rambutan. For each cell type, the difference in replication timing between each pair of loci in chromosome 21 was calculated. We then compared the average difference in replication time between highly ranked Rambutan predictions with high confidence Fit-Hi-C calls as a function of genomic distance (Fig. 6). We observe that highly ranked Rambutan calls are indeed indicative of shared replication timing. Furthermore, this effect only appears at distances greater than ~500 kb, as expected. Thus, Rambutan predictions are consistent with Hi-C contact maps in linking together synchronously replication regions of chromatin architecture.

**Figure 6:**
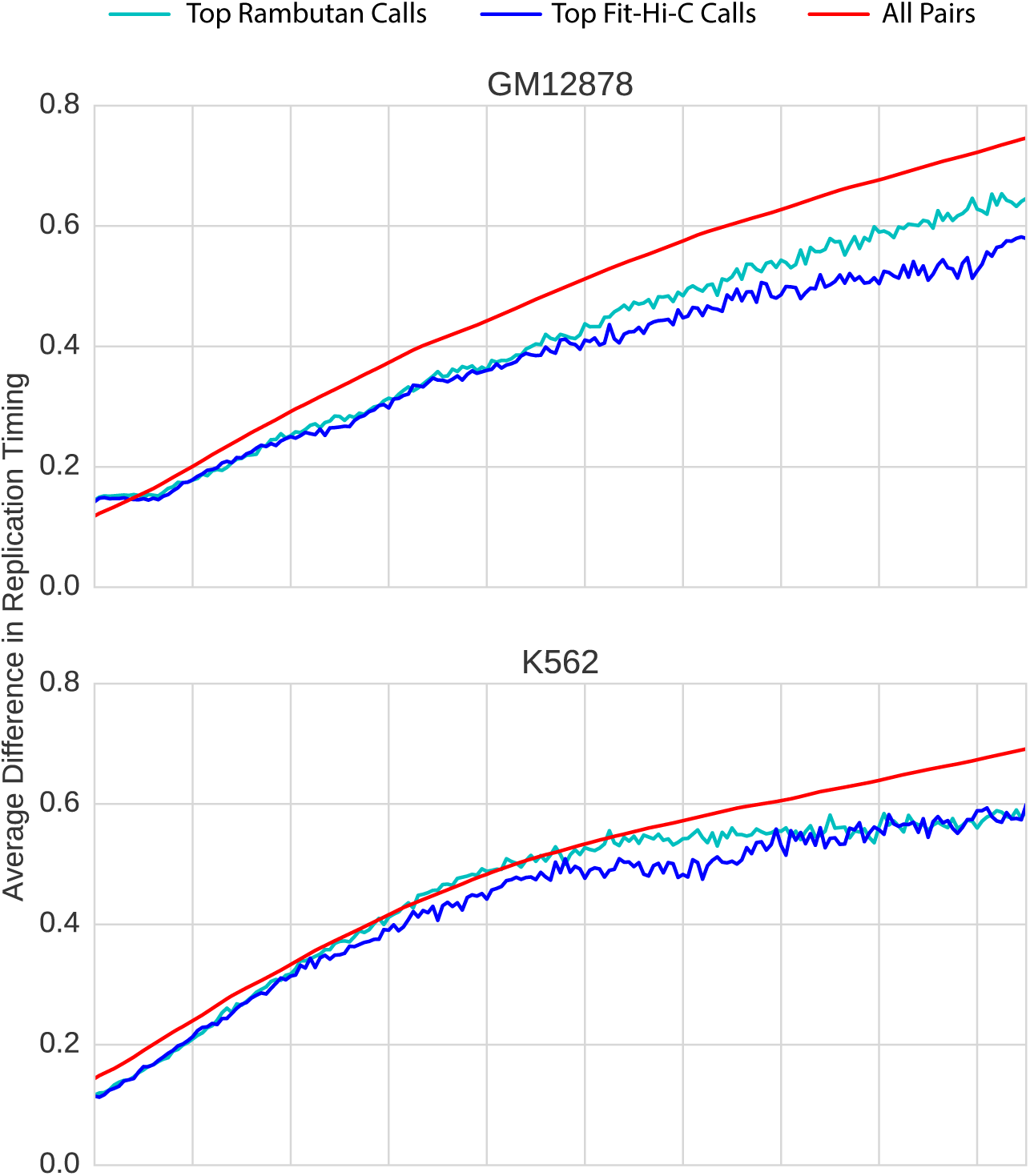
Differences in replication timing between loci in various sets. We calculate the mean difference in replication timing between loci in the top 1000 highest-confidence contacts according to Rambutan (cyan) and Fit-Hi-C (blue) at each genomic distance, as well as the mean difference in replicating timing across all contacts (red). The resulting values are shown for both Rambutan predictions and Fit-Hi-C p-values in cell types GM12878 and K562. The results for all cell types for which replication timing data is available are in the supplemental materials. Both methods show that high confidence predictions exhibit higher similarity in replication timing than other contacts, and that this divergence grows with respect to genomic distance.

### 4.5 Contact prediction for 53 human primary tissues

To demonstrate Rambutan’s utility in a fully predictive setting, we explored potential connections between cellular function and chromatin architecture, using data from the Roadmap Epigenomics Consortium. This data contains multiple epigenetic marks, including DNase sensitivity, for a large number of human cell types. Specifically, we used Rambutan to predict chromosome 21 contact maps in the 53 Roadmap cell types for which DNaseI data was readily available.

To visually inspect the results, we project the cell types into two dimensions using t-SNE (Van der Maaten and Hinton, 2008) (Fig. 7). The algorithm takes as input a matrix of pairwise distances between cell types. For this, we used 1 minus the correlation between insulation scores as a distance metric. It is immediately apparent that the cell types form distinct clusters which have functional similarity; for example, stem cells and immune cells both cluster together. In addition, four epithelial cell types, including HMEC and NHEK, cluster together towards the top, and three digestive cell types (gastric, pancreas, and small intestine) cluster together below that. Interestingly, a blood cell type and a leukemia appear close together, suggesting that possibly cancer has altered the structure detrimentally. In contrast, all cancer cell types (green dots) appear distant from another, likely due to cancer’s nature as a degradation of traditional cell function as opposed to a distinct functional class of its own.

**Figure 7:**
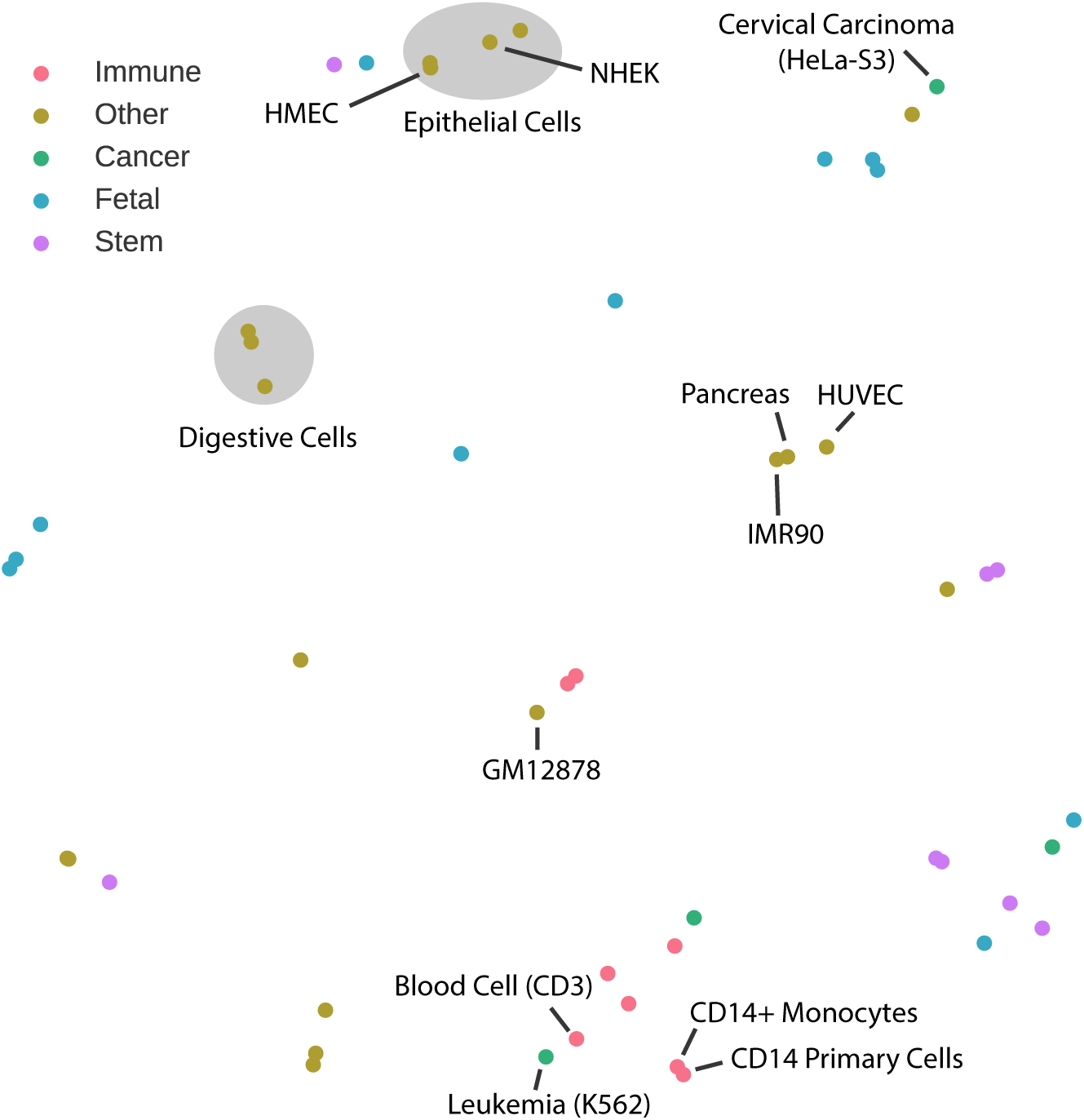
Visualization of a low dimensional embedding of cell type structures. The algorithm t-SNE was used to project cell types from a correlation based distance matrix into 2D space. A selection of five interesting cell groups are used to color the cell types with annotations derived from the Roadmap Epigenomics Consortium and the Regulatory Elements DB(Sheffield *et al*., 2013). All six of the originally inspected cell types are indicated on the plot, as well as various other cell types of interest. As examples, pancreas is included due to its proximity to both HUVEC and IMR90, and cervical carcinoma is included due to its distance from leukemia and the other cancer cell types.

## 5 Discussion

Rambutan is a deep neural network that predicts 3D chromatin architecture at 1 kb resolution using only nucleotide sequence and DNaseI signal. Notably, the model enables generation of predicted Hi-C contacts in cell types for which no Hi-C data exists. In addition to confirming that Rambutan makes accurate predictions within and between cell types, we provided three independent lines of reasoning to show that Rambutan’s predictions are consistent with known biology. First, we showed that insulation scores derived from Rambutan predictions strongly correlate with DNaseI peaks, replication timing, and several types of histone modifications. Second, we demonstrated that high confidence Rambutan predictions bring together loci with similar replication times. Third, we found that Rambutan predicts similar enrichment of contacts among several types of annotated gene regulatory elements and a depletion of contacts in quiescent regions. This result correlates strongly with enrichments calculated using experimentally derived Hi-C contact maps.

A major difficulty in the prediction of long-range Hi-C contacts is accounting for the strong genomic distance effect. Although the GM12878 data that we used to train the model contains many longer-range contacts (> 50 kb), the ratio of positives to negatives (high-confidence contacts versus not) is extremely skewed at higher distances. We used a balanced sampling scheme to ensure that the model receives a sufficient number of positively labeled examples. However, it is difficult to expose the model to enough contacts further than 1 Mb apart while not skewing the proportion of contacts seen at each genomic distance. A future direction may be to determine a training and validation scheme that accounts for this imbalance and allows for Rambutan to make more accurate predictions past 1Mb.

Another possible future direction is to train the model not as a classifier but as a regressor that predicts statistical confidence. This model could be more useful because it would allow users to set their own confidence thresholds. In addition, explicitly training the model to rank locus pairs would make more sense in contexts where the top *N* contacts are needed rather than contacts with confidence better than a specified threshold. A difficulty likely faced in this setting is that statistical confidence is tightly connected to how deeply sequenced that cell type is, so training data would have to be curated in order to account for this effect.

Recent advances in interpretable machine learning may soon allow complex models like Rambutan to provide succinct answers as to why they made a prediction in addition to the prediction. Most of these methods currently involve a sampling-based approach for explaining single predictions which identify which inputs were relevant in classifying that particular sample (Shrikumar *et al*., 2017; Ribeiro *et al*., 2016; Lundberg and Lee, 2017; Bach *et al*., 2015). A downside is that these techniques can be computationally intensive for models like Rambutan that require a large number of inputs.

Rambutan shows great promise for predicting significant contacts in all human cell types for which DNase data is available. Even for cell types without DNase data, acquiring DNase-seq data is far cheaper than performing a Hi-C experiment sequenced deeply enough to produce a 1 kb or 5 kb resolution contact map. Furthermore, a model similar to Rambutan might be produced by training from alternative chromatin accessibility assays such as ATAC-seq (Buenrostro *et al*., 2013), though we have not yet tried this. An interesting application of Rambutan would be to consider the role that genetic alterations might play in chromatin architecture and what connection may exist between these structural changes and disease. In particular, it may be possible to determine what differences in chromatin architecture are present between specific cancers and their respective healthy cells, leading to a more precise understanding of which treatments may be most effective.

